# A historical sequence deletion in a commonly used *Bacillus subtilis* chromosome integration vector generates undetected loss-of-function mutations

**DOI:** 10.1101/2024.01.04.574214

**Authors:** K. Julia Dierksheide, Gene-Wei Li

## Abstract

Since the 1980s, chromosome-integration vectors have been used as a core method of engineering *Bacillus subtilis*. One of the most frequently used vector backbones contains chromosomally derived regions that direct homologous recombination into the *amyE* locus. Here, we report a gap in the homology region inherited from the original *amyE* integration vector, leading to erroneous recombination in a subset of transformants and a loss-of-function mutation in the downstream gene. Internal to the homology arm that spans the 3′ portion of *amyE* and the downstream gene *ldh*, an unintentional 227-bp deletion generates two crossover events. The major event yields the intended genotype, but the minor event, occurring in ∼10% of colonies, results in a truncation of *ldh*, which encodes lactate dehydrogenase. Although both types of colonies test positive for *amyE* disruption by starch plating, the potential defect in fermentative metabolism may be left undetected and confound the results of subsequent experiments.

## Main text

The model Gram-positive bacterium *Bacillus subtilis* is widely used for strain engineering due to its natural competency and efficient homologous recombination system^1,2^. Synthetic DNA is commonly introduced into specific loci of the genome via homology-containing integration vectors that can be constructed and manipulated as plasmids in *Escherichia coli* (Figure 1A). One of the first genomic loci developed for integration vectors is at the gene *amyE*, which encodes α-amylase, a protein involved in starch degradation^3,4^. Successful integration leads to disruption of *amyE*, which can be easily screened for using an iodine stain that changes coloration upon binding to starch (“starch test”)^2^. The original *amyE* double-crossover integration vector pBGtrp and its derivatives, such as pDR111 and pDG1661^5,6^, have enabled studies on many aspects of microbiology, ranging from gene regulation to cell division^7–10^. They have also been central to the development of synthetic biology toolkits for *B. subtilis*^11–15^.

**Figure 1:**
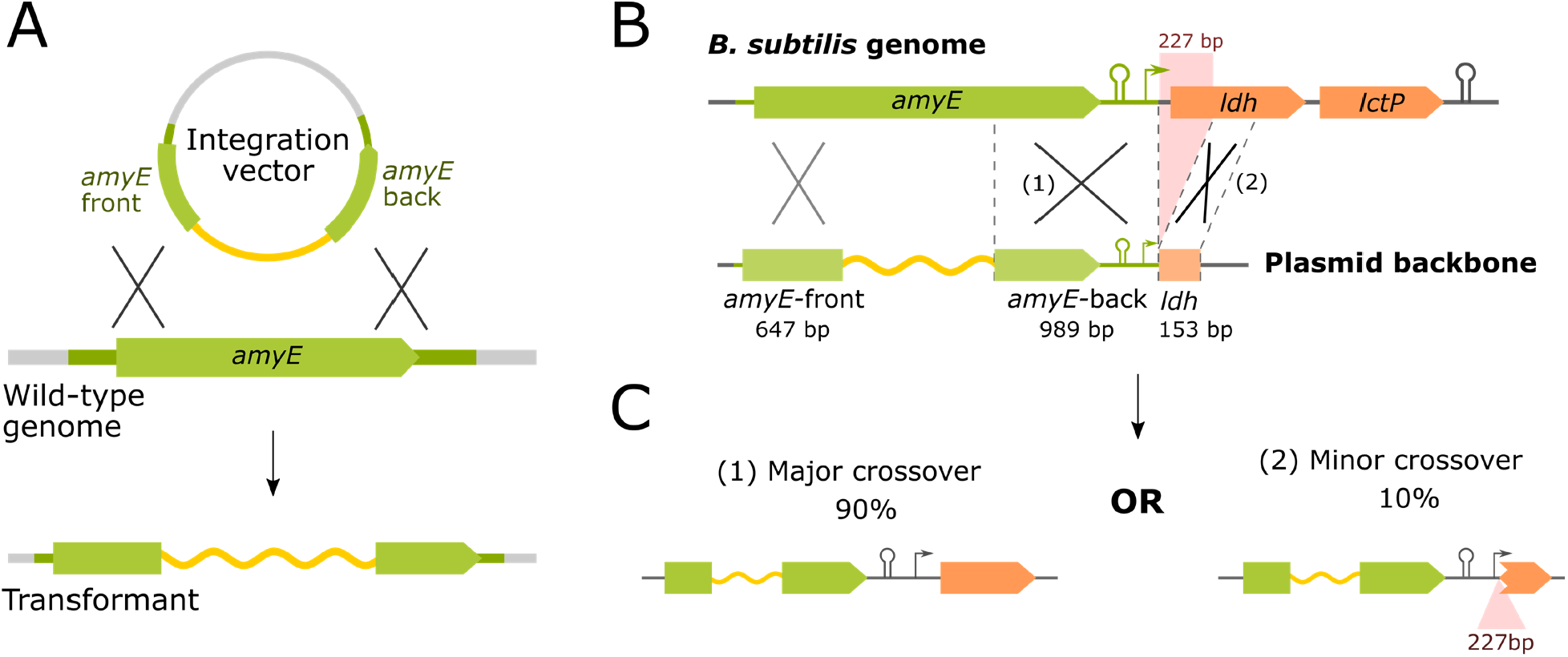
Double-crossover events at *amyE*. (A) Schematic of an *amyE* integration vector (top) designed to direct integration of the insert (yellow) into the genome as shown in the transformant genome (bottom). On the integration vector, the insert is flanked by two homology arms, *amyE*-front and *amyE*-back (green). (B) Schematic of the missing homology region. In the *B. subtilis* genome, *amyE* is followed by the *ldh-lctP* operon (top). In pBGtrp and its derivatives, the annotated *amyE*-back region is followed by a 153-bp fragment of *ldh*, while missing the intervening 227-bp sequence (bottom). (C) The two possible double-crossover events. In both cases, crossover occurs as expected at the upstream *amyE-*front region, but the missing genome sequence in the plasmid allows for two possible recombination events at the downstream *amyE-*back region. The minor event results in loss of 227 bp of genomic sequence containing the ribosome binding site and the first 215 nucleotides of *ldh*.

However, we found that the homology regions in these commonly used *amyE* integration vectors are inconsistent with the genome sequence^16^. In the sequence of pDR111, the annotated *amyE-*back homology region is followed by an additional 153-bp sequence derived from a region of the genome 227 bp downstream of *amyE-*back (Figure 1B). The resulting extended homology region includes a gap that belongs to the downstream *ldh* gene and its ribosome binding site. Due to this discontinuity in the homology region, in addition to the expected crossover at *amyE-*back, crossover can occur at the 153-bp region on the plasmid, disrupting *ldh*, a gene that codes for lactate dehydrogenase (LDH)^17^ (Figure 1C). We found that 4 of the 36 colonies tested after transformation with a derivative of pDR111 were missing the 227-bp region, indicating that the secondary crossover event occurs in a substantial proportion of transformants.

The discontinuous *amyE-*back homology region in pDR111 was inherited from pBGtrp, the original *amyE* double crossover integration vector developed in 1986^3,5,6,18,19^. The pBGtrp homology arms were generated from subclones of the *B. subtilis amyE* gene that were used to sequence the gene in 1983. We found that the corresponding sequence deposited in GenBank is missing the same 227 bp, indicating that this region was likely lost in the process of preparing *amyE* for sequencing in *E. coli*^4^. In addition to pDR111, many *amyE* double crossover integration vectors developed over the past 40 years, including pDG1661, likely have inherited the same discontinuous homology arms from pBGtrp and its derivative vectors.

To facilitate correction of this error in future work, we constructed modified plasmids of pDR110 and 111 where the 153-bp region downstream of *amyE-*back has been removed. The removal of the *ldh* homology region did not substantially impact transformation efficiency, and all colonies tested (18 of 18) integrated at *amyE* as expected for both plasmids. These plasmids will be available on AddGene as pGL003 (modified pDR110) and pGL004 (modified pDR111).

Historically, a single *B. subtilis* colony that tests positive by the starch test is carried forward after transformation for subsequent experiments. Our results suggest that, across all strains constructed with pBGtrp and its derivatives, ten percent of the strains may be missing the ribosome binding site and a major portion of LDH. Given LDH’s role in fermentative metabolism and anaerobic growth^20^, an undetected crossover in *ldh* may have influenced the results of previous experiments performed in these conditions.

This discrepancy can also influence studies with large-scale libraries of strains – whether pooled or arrayed – at the *amyE* site. Libraries of *B. subtilis* cells with pooled CRISPRi, overexpression, or reporter variants are powerful tools for discovery when coupled to modern high-throughput assays. When generating a library of *B. subtilis* variants, all cells that carry the intended antibiotic resistance cassette are carried forward from one or multiple transformation reactions. If the current, discontinuous *amyE* homology region is used, each transformed variant will integrate at *amyE* through one of the two possible crossover events (Figure 1C). These distinct crossover events are challenging to distinguish in high-throughput and introduce additional heterogeneity that could confound the results. Therefore, to ensure properly controlled experiments, especially in the context of fermentative *B. subtilis* studies, it will be important to correct the integration arms in future work.

## Acknowledgements

We thank A. D. Grossman’s laboratory for providing plasmids and members of the G.-W.L. and A.D.G. laboratories for discussions. This research was supported by NIH R35GM124732 and the NSF CAREER Award MCB-1844668.

## Supplementary Data

Sequence of pDR111 with annotations to indicate the position of *amyE-*front, *amyE-*back, and the homology region that includes the start of *ldh*.

